# Identification of the primary peptide inhibitor contaminant of fibrillation and toxicity in synthetic Amyloid-β42

**DOI:** 10.1101/108563

**Authors:** Daniel J. Adams, Travis Nemkov, John P. Mayer, William M. Old, Michael H. B. Stowell

## Abstract

Understanding the pathophysiology of Alzheimer disease has relied upon the use of amyloid peptides from a variety of sources, but most predominantly synthetic peptides produced using t-butyloxycarbonyl (Boc) or 9-fluorenylmethoxycarbonyl (Fmoc) chemistry. These synthetic methods can lead to minor impurities which can have profound effects on the biological activity of amyloid peptides. Here we used a combination of cytotoxicity assays, fibrillation assays and high resolution mass spectrometry (MS) to identify impurities in synthetic amyloid preparations that inhibit both cytotoxicity and aggregation. We identify the Aβ42Δ39 species as the major peptide contaminant responsible for limiting both cytotoxicity and fibrillation of the amyloid peptide. In addition, we demonstrate that the presence of this minor impurity inhibits the formation of a stable Aβ42 dimer observable by MS in very pure peptide samples. These results highlight the critical importance of purity and provenance of amyloid peptides in Alzheimer’s research in particular, and biological research in general.

## Introduction

Since introduction of the amyloid hypothesis of AD over 20 years ago[1], an overwhelmingly large literature has accumulated, cementing the central importance of Aβ42 in the mechanism of the disease [2,3,4,5,6,7]. Due to its causal role in AD, the Aβ42 peptide is a favored tool to model the disease mechanisms and phenotypes. Synthetically derived Aβ42 (sAβ42) has been widely used in animal models and cell culture systems to aid in understanding the biological targets and pathological mechanisms of AD. The sAβ42 is also heavily used for *in vitro* characterization of the basic biophysical properties which imbue Aβ42 with such unique and toxic potency. While the synthetic peptide has unquestionably proven to faithfully recapitulate much disease-relevant molecular and physiological pathology, data generated in these models notably suffer from poor reproducibility[8]. Because of its extreme propensity to aggregate in solution, harsh conditions are typically employed in the handling of Aβ42 to keep it soluble for use in various assays. These include concentrated urea, strong base, guanidine and even organic solvents such as hexafluro-2-isopropanol (HFIP). However, no single method is standard in the field. Seeding further variability, numerous conflicting methods are used in quantifying Aβ42 concentration, including ELISA, Absorbance 280, and BCA assays. Beyond these complications in handling Aβ, it is now becoming evident that the source and purity of the peptide can have major impacts on its performance in functional assays [8,9]. While sAβ42 is widely available from many manufacturers and has been used ubiquitously for many years, peptides produced via recombinant methods in bacteria are now emerging as an interesting alternative. Previous studies indicated that sAβ peptide contains impurities that alter its neurotoxicity and ability to aggregate[9]. Here we have validated those findings and extended the line of questioning to determine the identity of the contaminants which appear to inhibit the toxic activities of Aβ42. Through both discovery-oriented and candidate-based approaches, we have found that a failed valine-valine coupling at position 39-40 in Aβ42 produces a truncated peptide that co-purifies with full-length Aβ42 and is a potent inhibitor of its aggregation and cytotoxic activity. These results mandate that future assays using sAβ42 should ensure that this impurity is remove and we propose that this truncated Aβ derivative, and analogues thereof, merits further investigation in bioassays to characterize its potentially therapeutic properties against Alzheimer’s disease.

## Results

### Synthetic Aβ42 Exhibits Reduced Toxicity

Motivated by reports of divergent functional properties between recombinant and synthetic Aβ42[9], we sought to directly compare their cytotoxic potency. Samples of synthetic and recombinant Aβ42 were prepared as a monomer by solubilizing in hexafluoroisopropanol (HFIP) and re-drying to film under inert gas flow. The films were re-solubilized in 10 mM NaOH to prevent aggregation and diluted into culture medium immediately piror to application. The concentration of each peptide sample was determined by BCA assay in a method that has been independently verified by SDS-PAGE, UV-Vis spectroscopy and amino acid analysis. The popular pheochromocytoma-derived cell line PC12 was used in a toxicity assay because it is known to express many neuronal proteins and is easily amenable to high throughput bioassays. A concentration series of recombinant and synthetic peptide as well as the NaOH vehicle was prepared and applied to cells for 24 hours. At the endpoint, a standard MTT viability assay was performed (**Error! Reference source not found.**). We found that recombinant Aβ42 was potently toxic to the neuron-like PC12 cells, inducing measurable toxicity as low as 25 nM with an apparent LD_50_ of 190 nM. In contrast, the toxicity of the synthetic peptide was significantly lower, with an apparent LD_50_ of 280 nM.

### Synthetic Aβ42 Exhibits Qualitatively Different Fibrillation Characteristics

The aggregation behavior of Aβ42 is well documented and highly investigated, but the relationship between this behavior and the protein’s acute cytotoxicity is not entirely clear. Therefore, negative stain electron microscopy was used to determine if there were structural correlates of the reduced toxicity displayed by sAβ42. Accordingly, rAβ42 and sAβ42 stocks at 10 µM were diluted to 1 µM in phosphate buffered saline (PBS pH 7.4) to induce aggregation. Samples were removed for imaging after 60 minutes and analyzed by negative stain electron microscopy (**Error! Reference source not found.**). Both samples show aggregation into fibrils however the rAβ42 shows longer fibrils with little branching. In contrast the sAβ42 displayed shorter more branched clumps of fibrils

### Synthetic Aβ42 exhibits quantitatively different fibrillation dynamics

After observing qualitative differences in the morphology of aggregating fibrils of r Aβ42 and sAβ42, we sought to determine if differences might also exist in the quantitative dynamics of fibrillation. To this end we performed a standard fluorescence-based assay to track the kinetics of aggregation in real-time. Thioflavin T (ThT) is a benzothiazole dye that binds specifically to fibrillar Aβ and has been used for many years as a marker in histological identification of Aβ plaques in brain tissue. ThT binds an interaction surface unique to Aβ fibrils whereupon its fluorescence emission is greatly enhanced^109^. Thus an increase in ThT signal represents increased binding to fibrils and is therefore a direct readout for aggregation. Using the ThT dye assay, we observed distinct aggregation dynamics between rAβ42 and sAβ42 (**Error! Reference source not found.**). The kinetics of aggregation is set apart by two features of the fluorescent traces. Firstly, the lag time to reach the half-maximal signal, t_1/2_, is about 10% longer in the sAβ42 reaction indicating alterations to the early aggregation steps. Secondly, the fast-phase aggregation trajectory is less steep for the sAβ42. Together with the EM data, this indicates that the more highly branched fibrils observed in the synthetic sample form with reduced kinetics when compared to the more linear fibrils of recombinant peptide (**Error! Reference source not found.2**).

### Mutations within the Aβ42 Glycine Zipper Motif Alter Fibrillation and Toxicity

We have observed that the ability of Aβ42 to aggregate correlates with its toxicity. This correlation has been noted before[9,10,11], however it is not clear by what mechanism the tendency toward fibrillation would drive toxicity. To further explore this connection, we assayed if manipulations which impact aggregation behavior could also alter toxicity. For this experiment we utilized the G37L mutant of the Aβ42 peptide. This mutation disrupts the glycine zipper motif (**Fig 4**) that is important for the normal homo-oligomerization of Aβ42[10,12]. It has been observed that the G37L mutant peptide can act as a dominant negative in an aggregation assay[9,10,13]. Thus, we performed a 1:1 mixing experiment between wild-type and G37L Aβ42 and applied this peptide to PC12 cells in the same toxicity experiment as above (**Error! Reference source not found.**). We found that the G37L peptide exhibited a protective effect from Aβ42 toxicity (**Fig 5**). This result supports the claim that features of Aβ42 involved in oligomerization are also important in toxicity.

**Fig 1:**
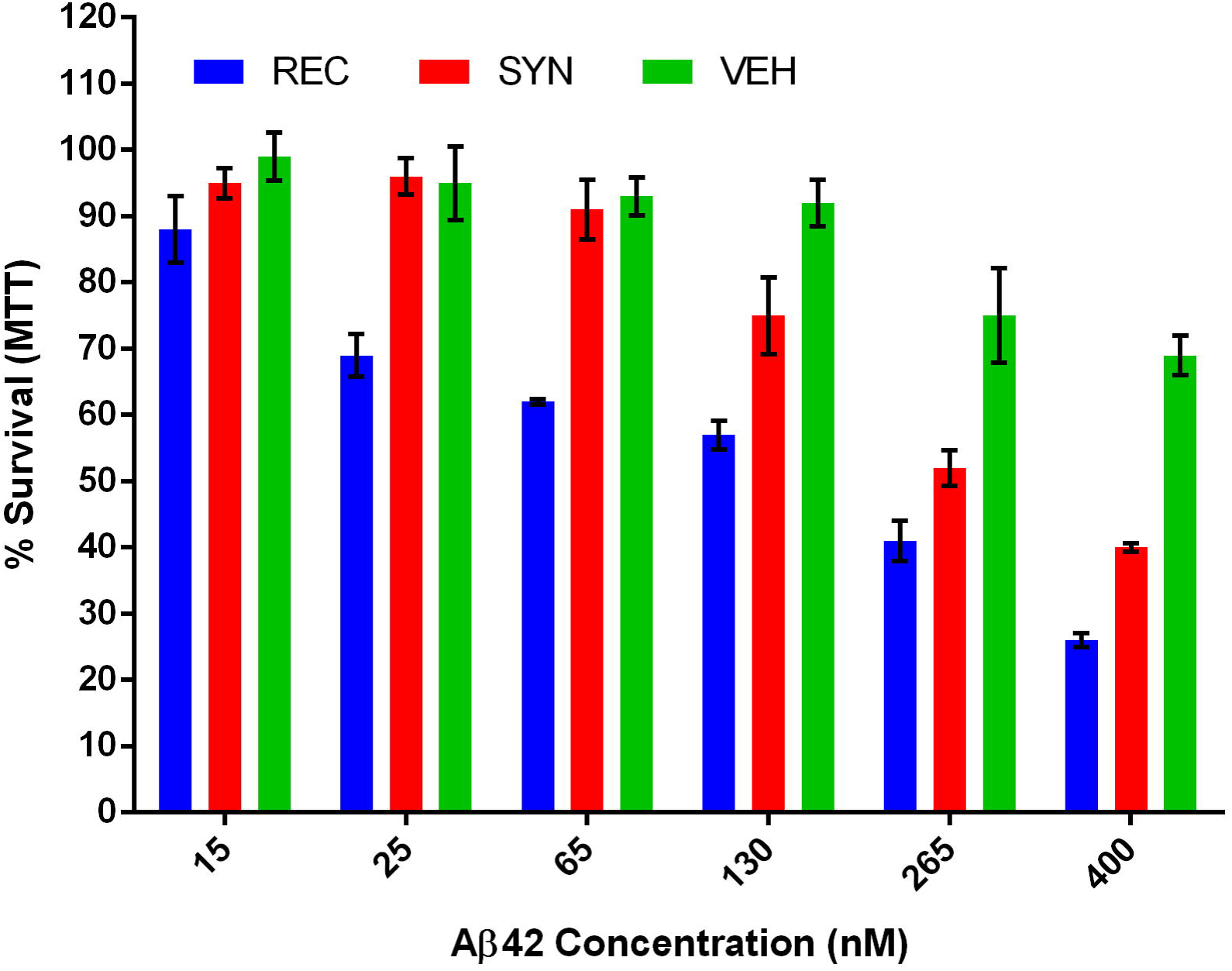
Recombinant Aβ42 is more cytotoxic than synthetic Aβ42. MTT survival assay after 24h of peptide treatment at indicated concentrations. Averages from two experiments and six replicates. Error bars represent SEM. A pairwise T-test across all dose series gave a p=0.004 comparing the synthetic (SYN) and recombinant (REC),a p=0.001 comparting REC and vehicle (VEH), and a p=0.06 comparing SYN to VEH.

**Fig 2:**
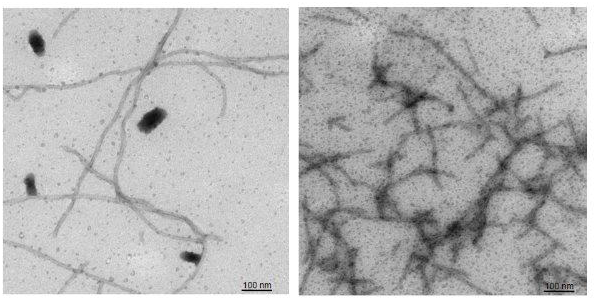
Aβ42 displays more uniform fibril formation than synthetic Aβ42. Negative stain EM analysis of recombinant Aβ42 (left) versus synthetic Aβ42 (right).

**Fig 3:**
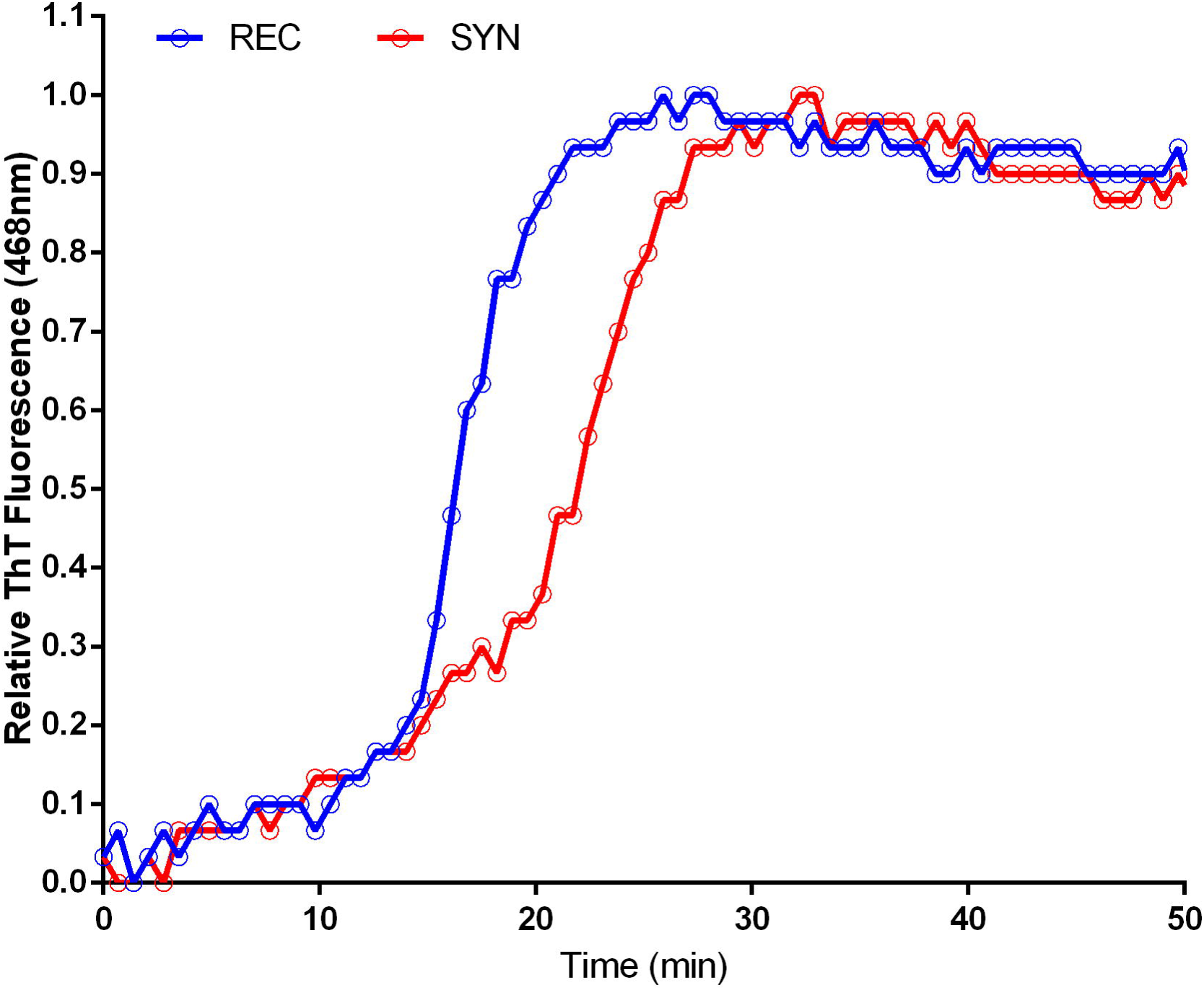
Aggregation of Recombinant, Synthetic and Mixed Aβ42. ThT fluorescence trace of Aβ42 aggregation. REC is recombinant and SYN is synthetic Aβ42.

**Fig 4:**
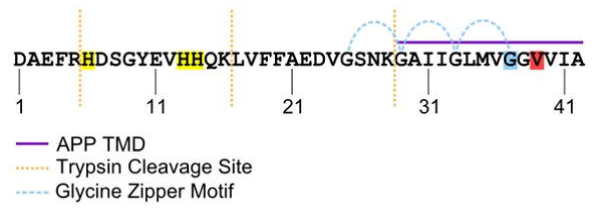
Sequence and Features of Human Aβ42. Sequence of human Aβ42 peptide. Histidine residues highlighted in yellow. Glycine 37 highlighted in blue. Valine 39 highlighted in red. Purple bar indicates sequence derived from the transmembrane domain of APP. Blue curved lines indicated glycine zipper motif.

**Fig 5:**
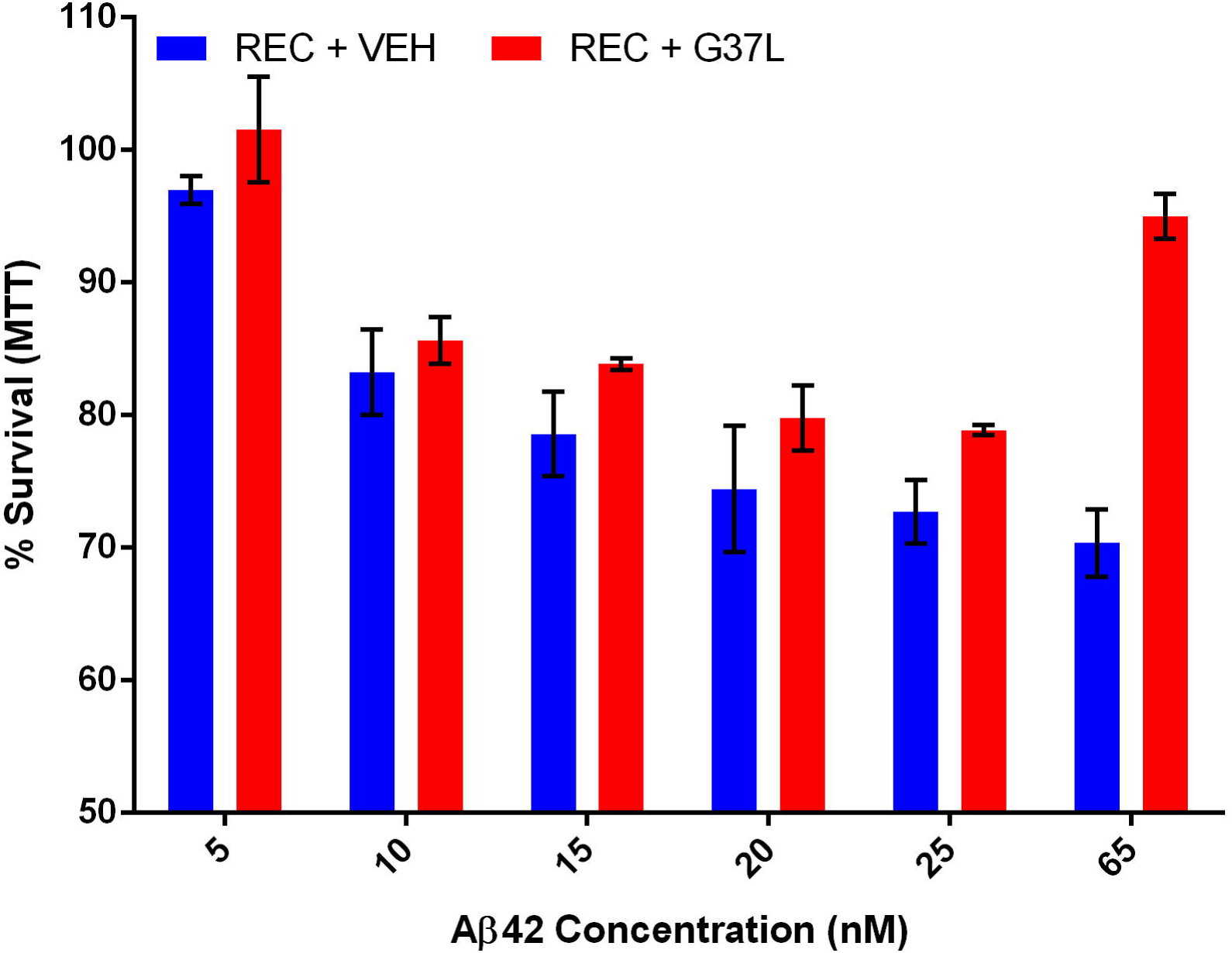
Aβ42 G37L Protects from Toxicity as Dominant Negative. Wild-type recombinant Aβ42 was mixed in an equimolar ratio with Aβ42 G37L. PC12 cells were treated for 24 hours and viability was assayed by MTT. Bars represent 6 replicates. Error bars represent SEM. A pairwise T-test across all dose series gave a p=0.06 comparing the recombinant (REC) and recombinant doped with the G37L mutant (REC + G37L).

From multiple manufacturers, a typical lot of commercial Aβ42 peptide is stated as >95% pure. Assuming these numbers are accurate, two distinct scenarios could explain the disparity in aggregation dynamics and toxicity between recombinant and synthetic Aβ42. One possibility is that some contaminant in the recombinant peptide is responsible for its comparatively enhanced toxicity and aggregation behavior. If it were possible to identify this contaminant and add it to the synthetic Aβ42, increased performance of the synthetic peptide would be expected. The other possibility is that a minority product present in the synthetic peptide is inhibiting its natural aggregation and toxicity. To distinguish between these possibilities, a doping experiment was performed in which a small amount (5% mole fraction) of synthetic peptide was added to the recombinant peptide and subjected to aggregation analysis by ThT fluorescence. If a contaminant of the recombinant peptide were responsible for its enhanced aggregation behavior compared to the synthetic peptide, then no change in dynamics would be expected; but if an inhibitor of aggregation were present in the synthetic peptide, it should still be able to produce a measurable effect even diluted 20 fold. Strikingly, we observed that only 5% of the synthetic peptide was sufficient to perturb the normal dynamics of Aβ42 aggregation (**3**). Both the time to t_1/2_ and slope of rapid aggregation phase indicate that the dynamics of aggregation have indeed been altered by the minority species present in the synthetic peptide. In contrast, 5% doping of recombinant Aβ42 into the synthetic peptide was not able to improve aggregation behavior (**Error! Reference source not found.**), again suggesting the sAβ42 contains an inhibitor of aggregation. We next used this doping strategy to assess toxicity of Aβ42 mixtures in PC12 cells. In agreement with our ThT data, just a small dose of synthetic peptide was capable of reducing the toxicity of the recombinant peptide by a measurable amount (**Fig)**, further confirming the potency of the inhibitory species present in the synthetic Aβ42 preparation.

### Identification of Aβ Inhibitors

Our data indicate that a potent inhibitor of both the aggregation (**Error! Reference source not found.**) and toxicity (**Fig 6**) of Aβ42 exists as a minority product present in the commercially obtained synthetic peptide. This finding has wide implications due to the ubiquitous use of synthetic Aβ42 in AD research. However, due to the recent commercial availability of recombinant Aβ42, it may not be necessary to improve upon commercial synthesis and purification to avoid these inhibitory contaminants. Instead, these unknown inhibitors may prove valuable as a starting point for the rational design of molecules for therapeutic intervention in AD. Furthermore, these inhibitors serve as a unique and novel tool for probing the relationship between Aβ42 aggregation and toxicity.

**Fig 6:**
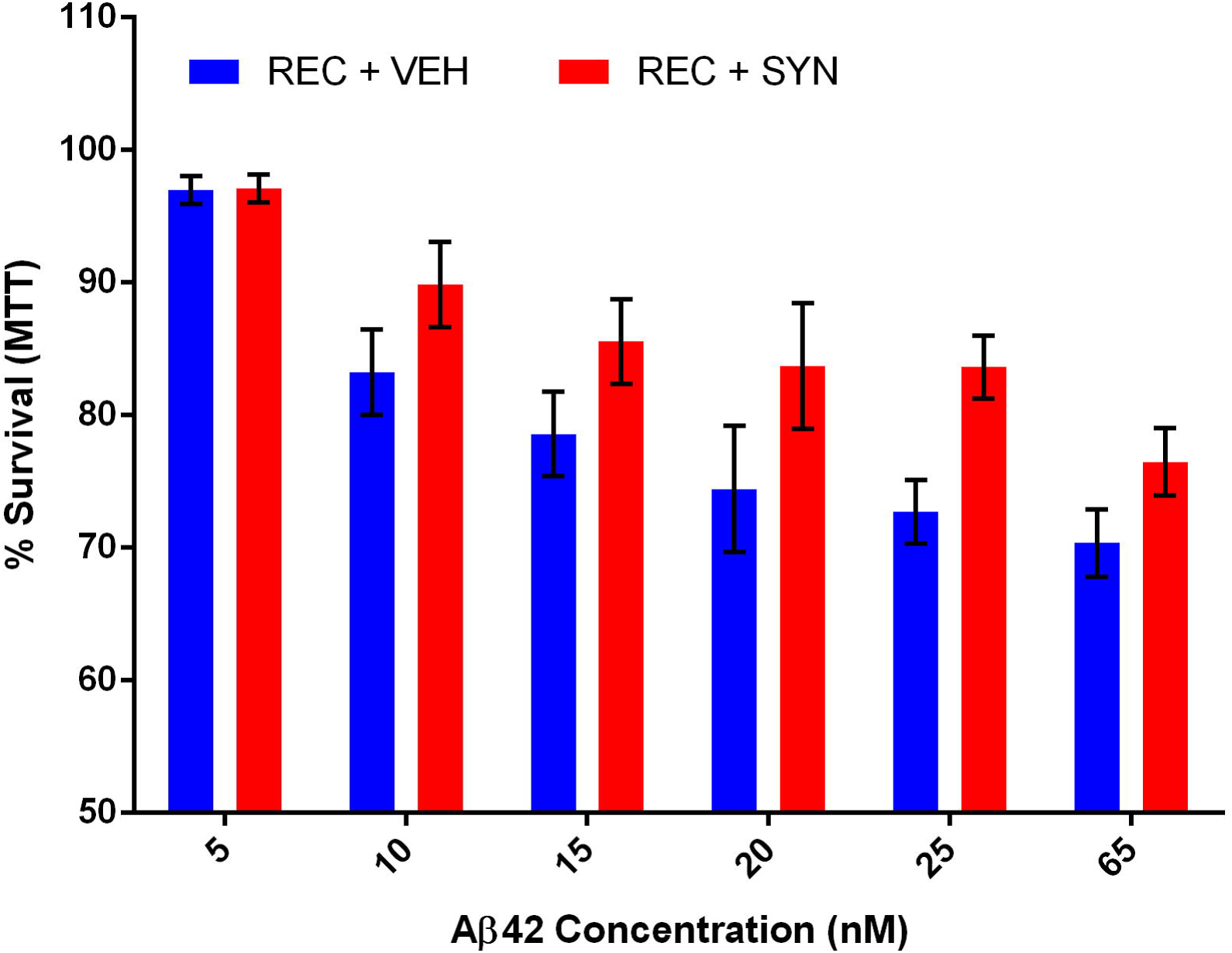
Aβ42 Contains a Potent Inhibitor of Toxicity. MTT toxicity assay of PC12 cells treated with recombinant Aβ42 ± 5% synthetic Aβ42. Assay performed in triplicate, error bars represent SEM. A pairwise T-test across all dose series gave a p=0.007 comparing the recombinant (REC) and recombinant doped with 5% synthetic (REC + SYN).

We began our study of the contaminants of synthetic Aβ42 by optimizing a chromatographic method for both purification and analysis. We found conditions under which we could elute peaks of recombinant Aβ42 from a C8 reverse-phase column in a mobile phase of acetonitrile with TFA as a counter ion. We then used this method to analyze the synthetic Aβ42 sample for the presence of contaminants. The same sharp main peak existed, however a number of additional minor peaks were observed. There was a distinct peak that eluted just before the main peak, as well a significant shoulder on both the leading and trailing edges of the main peak (**Fig 7**).

**Fig 7:**
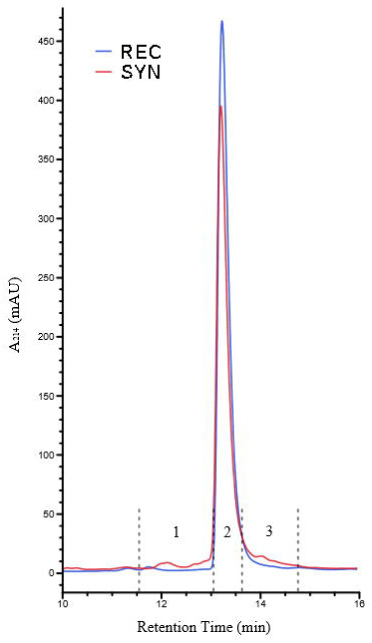
HPLC Analysis of Recombinant and Synthetic Aβ42. RP-HPLC of recombinant and synthetic Aβ42 reveals contaminants in synthetic preparation. 10 µg of each peptide sample was loaded. Fractions collected from synthetic sample are indicated 1, 2 and 3 by dashed lines.

We also analyzed both recombinant and synthetic peptide samples by MALDI-TOF mass spectrometry (**Fig 8**). This analysis revealed that the synthetic peptide sample contains a diverse collection of contaminants; however due to their relatively low abundance no specific identifications from this MALDI-TOF cocktail were possible. Most of the contaminants appear within approximately 400 mass units of the main peak, corroborating previous suggestions [6] that they may be byproducts of synthesis related to the majority product. A loss or addition of one to three residues would result in truncated or augmented peptides consistent with the observed sizes of contaminants.

**Fig 8:**
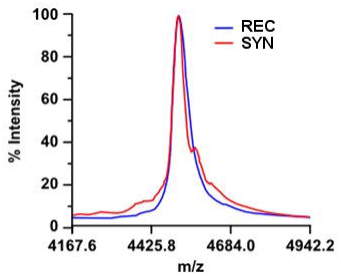
Mass Spectra of Aβ42 Samples. MALDI-TOF spectra reveal major contaminants in synthetic Aβ42 sample.

It has been reported that synthetic Aβ42 contains a significant fraction (>1%) of D-histidine that is suggested to be the functionally relevant contaminant in the synthetic material[6]. The traditional reverse-phase chromatographic methods used to purify Aβ42 would be incapable of resolving peptides epimerized at any of the three histidine residues in Aβ42. Racemized peptide would also be indistinguishable by mass spectrometry, making it very difficult to assay for this contaminant with commonly used tools. To directly test if an Aβ42 histidine racemate was capable of recapitulating the aggregation inhibition imparted by the synthetic peptide, we obtained a sample of racemized peptide for ThT assay. The peptide was synthesized using a standard solid phase Fmoc-protected method, but at positions 6, 13 and 14 a 50:50 mixture of L- and D-histidine enantiomers was applied. Thus, a mixture of Aβ42 molecules representing every possible combination of histidine stereochemistry was yielded (noted below as Aβ42-HIS). The Aβ42-HIS was doped into recombinant Aβ42 at 5% mole fraction, a much higher concentration than might normally appear in the synthetic peptide, and aggregation was monitored by ThT fluorescence (**Fig 9**). We found that even this large dose of Aβ42-HIS produced a minor alteration in the recombinant peptide’s velocity of aggregation. Therefore, we conclude that although Aβ42-HIS may be capable of inhibiting Aβ42 aggregation, it alone is not sufficient to illicit effects of the magnitude observed. Therefore, we believe that other as yet undiscovered inhibitors in synthetic Aβ42 must exist.

**Fig 9:**
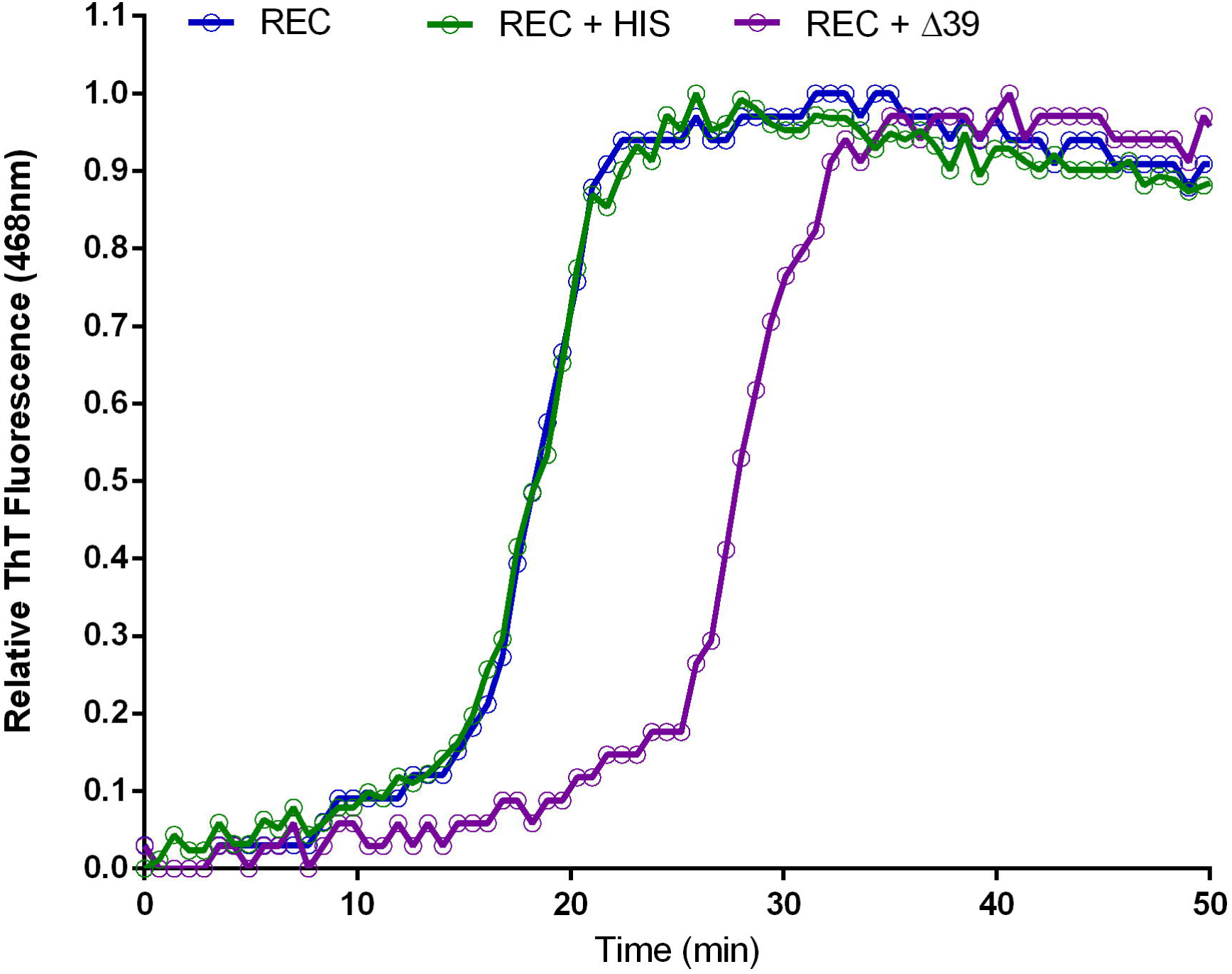
Aβ42 Mutation Aggregation Kinetics. ThT fluorescence was used to assay aggregation dynamics of 5% Aβ42-HIS mixed into recombinant Aβ42 (REC + HIS) or 5% Aβ42Δ39 (REC + Δ39).

Using RP-HPLC (**Fig 7**) and mass spectrometry (**Fig 8**), we observed many contaminating minority products in synthetic Aβ42. To determine if any of these possessed inhibitory activity, synthetic Aβ42 samples were fractionated by RP-HPLC and then assayed for inhibition of aggregation using the ThT assay. Specifically, the main peak of synthetic Aβ42 was isolated from the contaminating material that eluted on either side of it, yielding fractions labeled 1, 2 and 3 (**Fig 7**). As before, fractions were doped into recombinant Aβ42 at 5% wt/wt for the aggregation assay. Significant changes in aggregation dynamics were not observed in recombinant peptide doped with fractions 2 and 3, but we found that fraction 1 contained extremely potent inhibitory activity (**Fig 10**). It is important to note that this fraction was still a complex mixture; therefore, it remains unclear whether its inhibitory activity is caused by a single or multiple species. Without further analysis by more sensitive methods such as tandem MS-MS, it is impossible to ascertain the identity of the active agents isolated in fraction 1, although based on the MALDI-TOF experimental results (**Fig 8**) we suspect them to be related peptides. With the quantity of relevant contaminants available for study being extremely limiting we were prompted us to adopt a candidate-based approach towards identification of Aβ42 activity inhibitors.

**Fig 10:**
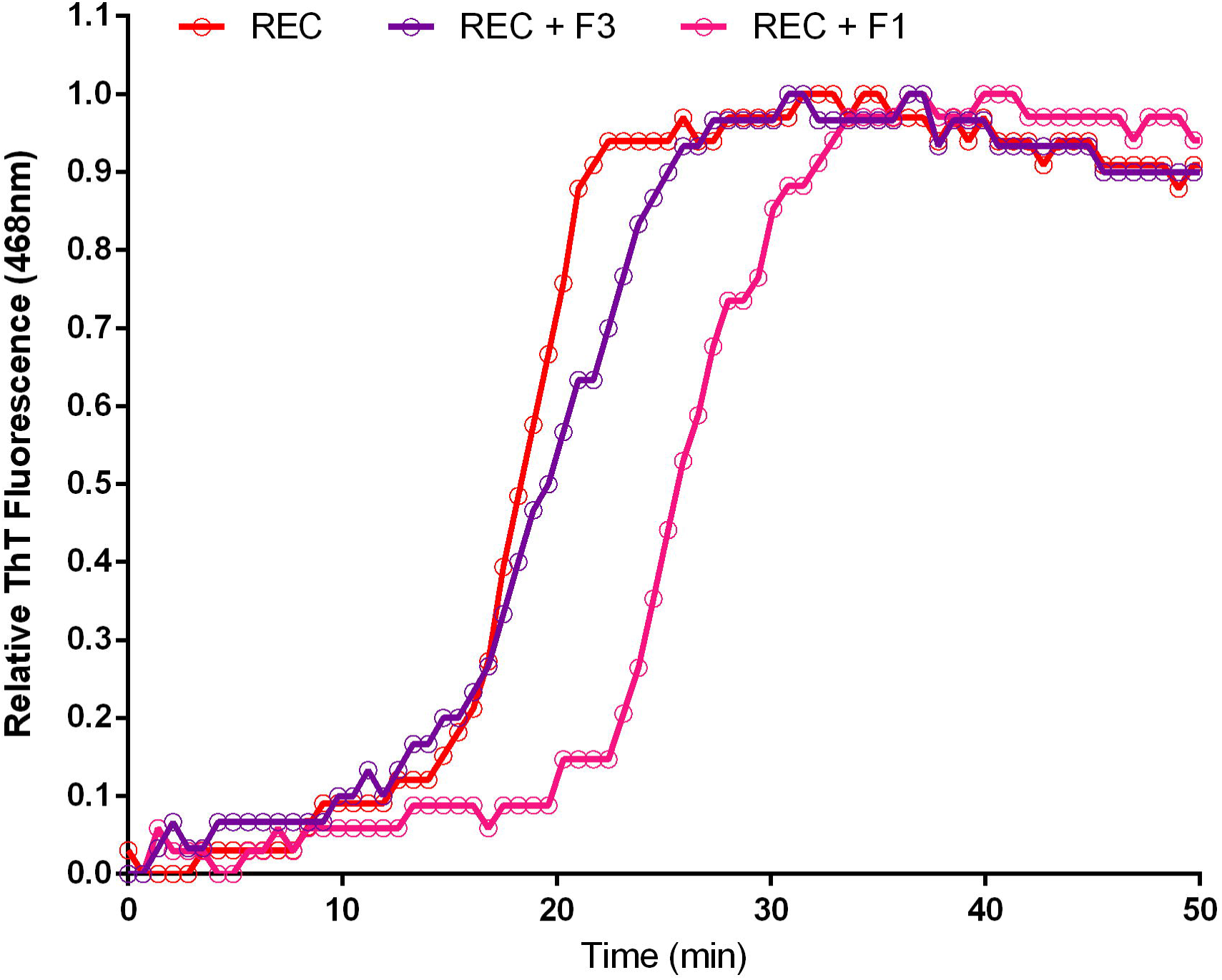
Contaminant is Potent Inhibitor of Aggregation. HPLC fraction 1 and 3 was doped into recombinant Aβ42 at 5% wt/wt and aggregation was assessed by ThT fluorescence. Fraction 1 contained a potent inhibitor of Aβ42 aggregation.

There are several sequence features which are known to pose challenges to standard solid phase peptide synthesis methods, and therefore can inform rational prediction of peptide synthesis byproducts for a given target. Highly hydrophobic aggregation-prone sequences such as found in Aβ42 are known to be particularly difficult to generate[14]. Traditional synthesis methods proceed from the carboxy-terminus towards the amino-terminus coupling a single amino acid at a time each followed by a round of deprotection. Given that the carboxy-terminal ^1^/_3_ of Aβ42 is derived from the transmembrane helix of APP (**Fig 4**), early steps in the synthesis of Aβ42 are particularly susceptible to aggregation of the nascent peptide chain[14]. Peptide aggregation competes with binding of synthesis reagents including deprotection agents and subsequent amino acid residues. Because of the iterative nature of the synthesis process, if either deprotection or coupling fails for a particular molecule in one round, that molecule may well participate in future rounds of synthesis, yielding a final peptide lacking only a single amino acid. In addition to these challenges, valine to valine coupling reactions, which are used in Aβ42 synthesis, are known to be of lower efficiency[15] than other peptide couplings in general. These facts led us to predict that omission of valine 39 (hereinafter Aβ42Δ39) generated a putative byproduct of synthesis that could be responsible for the observed inhibitory activity. This hypothesis is supported by the presence of a peak mass observed by MALDI mass spectrometry approximately 100 Da below the main peak. Additionally, the HPLC fraction containing the inhibitory activity eluted slightly before Aβ42, indicating slightly reduced hydrophobicity that is also consistent with deletion of a valine residue.

Absolute mass resolution by MALDI-TOF is inversely correlated with analyte mass[16], and therefore the 3% difference in mass of Aβ42Δ39 from the total peptide would prove difficult to resolve using solely this method. Digestion of the sample with trypsin would generate a C-terminal fragment Aβ42 29-42 (**Fig 4**). Aβ42 29-42 has an expected mass of approximately 1270 Da which would shift to 1171 Da upon deletion of Val 39, a change of nearly 10% which would be more readily resolved. Therefore, this combinatorial method of enzymatic digestion and subsequent MALDI-TOF analysis was performed in order to reveal putative Aβ42Δ39 contaminants. Digestion was carried out on synthetic Aβ42 overnight at 37 °C. The resultant peptides were desalted/concentrated with a C_4_ reversed-phase ZipTip and eluted in acetonitrile and analyzed by MALDI-TOF.

After significant optimization of desalting conditions and matrix/spotting parameters we were able to identify the expected four trypsin digestion peaks for fragment AA1-5, observed mass 638.21 (Calc. 637.29); fragment AA6-16, observed mass 1337.23 (Calc 1337.60); fragment AA17-28, observed mass 1324.79 (Calc. 1325.67); and fragment AA29-42, observed mass 1271.55 (Calc. 1269.76). We did not observe any peaks corresponding to partial digestion indicating that the reaction had gone to completion. Disappointingly, the 29-42 peptide of interest, containing Aβ42Δ39, was found to ionize very poorly by MALDI, leading to very weak signal. We did not see evidence of a peak near 1153Da although we considered this uninformative as the Aβ42Δ39 would represent only a very small fraction of the already weak signal from the 29-42 fragment.

To overcome the sensitivity hurdles of MALDI-TOF in the identification of potential contaminants, we turned to ESI-Orbitrap mass spectrometry analysis on both recombinant and synthetic Aβ42 (**Fig 11**). This data confirmed the presence of the many contaminating species in synthetic Aβ42 observed by MALDI MS and HPLC. The ESI-Orbitrap data allowed us to resolve individual peaks out of the shoulder present on the Aβ42 peak seen by MALDI MS from approximately 4200 Da to 4500 Da. Importantly, a peak of 4417.25 Da was observed, consistent with the presence of Aβ42Δ39. Based on this finding, we proceeded with direct evaluation of Aβ42Δ39 as an inhibitor of Aβ42 aggregation. Also of note was the presence of a peak in the recombinant sample of Aβ42 dimer, M/Z 1290.52. This indicates not only that small oligomers can form under the inhibitory conditions used (1% NH_4_OH), but that these oligomers can remain intact through electrospray ionization, trapping and detection. The absence of this peak in the synthetic sample appears to serve as further evidence of its reduced ability to aggregate.

**Fig 11:**
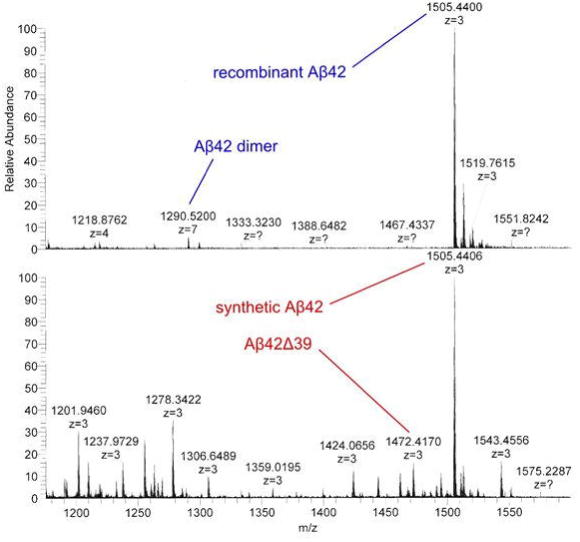
ESI-Orbitrap MS of Synthetic and Recombinant Aβ42. Top panel, ESI-Orbitrap MS analysis of recombinant Aβ42 show the presence of a dimeric species at 1290.52 M/Z and the lack of the Aβ42Δ39 peptide at 1472.42 M/Z. Bottom panel, ESI-Orbitrap MS analysis of synthetic Aβ42 showing the clear presence of the Aβ42Δ39 peptide at 1472.42 M/Z and many other contaminating species but undetectable levels of the 1290.52 M/Z dimer species.

Finally, to directly test if Aβ42Δ39 was capable of inhibiting Aβ42 aggregation kinetics, we obtained the modified peptide, mixed it into recombinant Aβ42 at 5% mole fraction, and assayed aggregation by ThT fluorescence. We observed a remarkable inhibition with a near two-fold increase of 
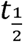
 from 17.9 minutes (Aβ42) to 30.5 minutes (Aβ42Δ39) (**Fig 9)**. This shift was reminiscent of the effect imparted by the HPLC fraction 1 (**Fig 10**) further corroborating that Aβ42Δ39 is the major active component of that fraction.

## Discussion

We have found that synthetic manufacturing of Aβ42 generates contaminating byproducts that inhibit Aβ42 aggregation. We have also identified Aβ42Δ39 as a potent agent of this inhibition. To a lesser extent, another suggested contaminant, Aβ42-HIS, also exhibited some inhibition of aggregation. Previously observed variably in assays utilizing Aβ42 can most readily be ascribed to the primary contaminants observed in synthetically prepared Aβ42. Although Aβ42 is particularly prone to synthetic contaminants arising from coupling inefficiencies, the potential for minor impurities to alter both the biophysical and biological properties of synthetically prepared peptides should be born in mind regardless of the peptide being investigated. Finally, and most intriguingly, the observed correlation between Aβ42 toxicity and aggregation kinetics, suggests that Aβ42Δ39 and perhaps Aβ42-HIS may exhibit neuroprotective activity in AD models. Future, cell-based assays will reveal whether treatment with Aβ42Δ39 or Aβ42-HIS can serve to attenuate Aβ42 toxicity, possibly yielding an important new class of compounds for therapeutic intervention in Alzheimer’s disease.

## Materials and Methods

### Materials

Thioflavin T was purchased from Sigma-Aldrich. Recombinant Aβ42 was obtained from either AmideBio or rPeptide. Synthetic Aβ42 was purchased from Bachem and Anaspec. Aβ42Δ39 was purchased from Anaspec. All other chemicals were of analytical grade.

### HPLC analysis

Samples were analyzed on an Agilent 1100 and fractions isolated using a 4.6 × 250 mm reversed phase Vydac MS C8 column (Grace) using a two phase elution comprised of 1% ACN + 0.1% trifluoroacetic acid (Buffer A) and 100% ACN + 0.1% trifluoroacetic acid (Buffer B) at 70°C. The elution profile was, 0–10 min; 0% B, 10–30 min linear increase to 80% B, with a constant flow rate of 0.8 ml/min and protein was detected at 215 nm.

### Electron microscopy

Samples for EM were prepared according similar to the methods of Komatsu[17]. Briefly, a 2% w/v aqueous solution of uranyl acetate at pH 4.2 was filtered (0.2 μm) to remove small precipitates and stored in foil covered container prior to use. Approximatly 10 ng (3 μL) of amyloid sample was place on a freshly glow discharged carbon film on a 300 mesh copper grid for 1 minute. The sample was blotted and then the uranyl acetate solution was applied for 30 seconds and then blotted and air dried. Images were recorded on side mounted CCD camera using a Phillips CM100 at 80 kV.

### ESI-Orbitrap Mass Spectrometry

Amyloid Beta samples were reconstituted in pH 10 1% NH_4_OH at 1 mg/mL. Samples were infused at 300 nl/min for nano-electrospray ionization and mass spectrometry analysis performed on a LTQ-Orbitrap (ThermoFisher). Survey scans were collected in the Orbitrap at 60,000 resolution (at m/z=300). The maximum injection time for MS survey scans was 500 ms with 1 microscan and AGC= 1x10^6^. For LTQ MS/MS scans, maximum injection time for survey scans was 250 ms with 1 microscan and AGC= 1×10^6^. Peptides were fragmented by CAD for 30 ms in 1 mTorr of N_2_ with a normalized collision energy of 35% and activation Q=0.25.

